# Genome-wide overexpression screen of *Pseudomonas putida* phage Emajogi

**DOI:** 10.64898/2026.09.15.751885

**Authors:** Ben Diaz, Stephen Won, Meghana Padala, Catherine M. Mageeney

## Abstract

Bacteriophage genomes contain large fractions of uncharacterized genes, motivating high-throughput functional screens to illuminate their roles in host interaction and cytotoxicity. Here, we performed a genome-wide overexpression assay for a *Pseudomonas putida* KT2440-infecting phage, Emajogi, and identified eight cytotoxic genes that reduce host growth under both basal and induced expression. We show that these proteins are conserved across phages that infect a diverse set of hosts. The most cytotoxic gene identified was a hypothetical protein (*gp18*) with no predicted functional domains. Transcriptomic profiling revealed that overexpression of gp18 caused a pronounced metabolic remodeling of the cell, likely shifting cell resources to phage production. Overall, this genome wide screen provides information about phage genes that were previously unannotated that can support biotechnology and novel antimicrobial development.

## Introduction

Bacteriophages are abundant, diverse, and present everywhere bacteria exist^1^. Bacteria and phages have been co-evolving for billions of years which has led to highly timed and complex processes for phage infection, which involves DNA injection, overcoming bacterial defense mechanisms, overtaking the bacterial cell to turn it into a phage factory, and lysis of the bacterial host^2^. The phage genome encodes genes that perform many of these processes; however, ascribing specific genes to these functions is limited. While structural proteins are well annotated due to obvious phenotypes upon removal, other functions including anti-defense, host takeover, and auxiliary metabolic genes are not as easily characterized.

Overexpression studies can help provide necessary insights to understand phage genomic “dark matter”^3^. Previous overexpression studies of bacteriophage genes have revealed novel mechanisms of integration, new anti-defense genes, genes involved in host take over, and genes involved in metabolic reprogramming^4-8^. Further, overexpression studies also provide information to develop additional assays to determine specific functions. Lastly, genes identified as cytotoxic may provide a defense against additional incoming phages or may lead to advances in antimicrobials and synthetic biology.

Here we report results of a genome-wide overexpression screen for Emajogi, a phage which infects *Pseudomonas putida* KT2440, a rapidly growing biomanufacturing chassis^9,10^ and prominent soil community member^11^. Emajogi is predicted to encode 50 genes with 18 genes having no assigned function (**Figure 1a**)^12^. Out of the individual 47 genes we cloned and tested, we found eight genes in Emajogi (17%) to be cytotoxic to *P. putida* KT2440. Two of these genes are annotated to encode hypothetical proteins, while the remaining six have roles in DNA replication and transcription or are structural proteins. We used transcriptomics to understand how overexpression of gp18, a cytotoxic hypothetical protein, impacts *P. putida*. The data suggests that gp18 is involved in host take over since operons related to oxidative stress and cell remodeling were differentially regulated.

**Figure 1.**
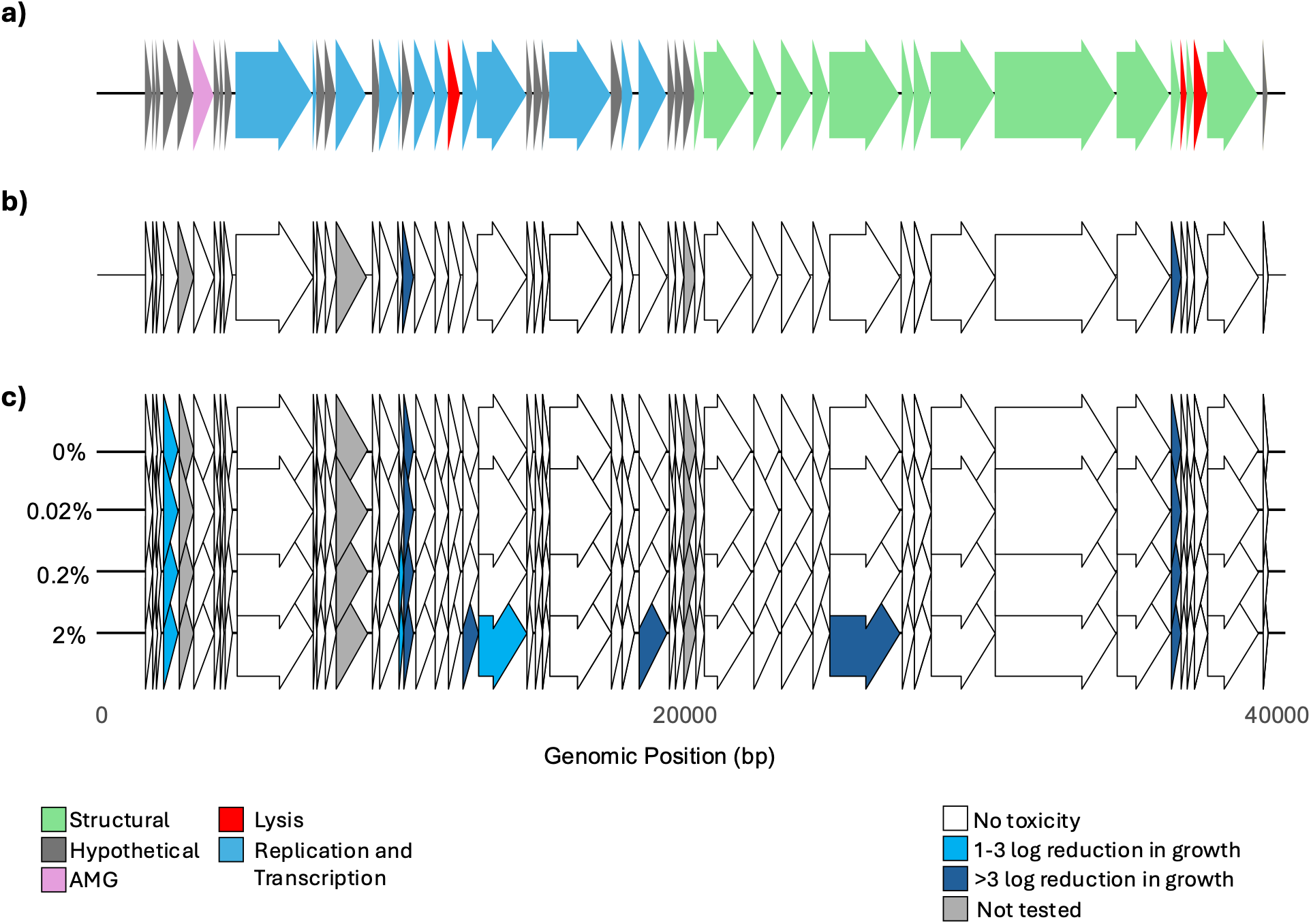
Inducible expression identifies cytotoxic gene products across *Pseudomonas putida* phage Emajogi. **a)** Gene map of Emajogi (NCBI accession number PP4946445.1), functional categories are denoted by color key on the lower right. **b**) Cytotoxicity scores in LB media. Because inducer concentration did not change the cytotoxicity scores, all are shown in a single map. **c**) Cytotoxicity scores in M9 media. Different concentrations of arabinose added are shown to the left of the genome. The color key representing the score or genes not tested is shown on the lower left for panels **b** and **c**.

## Results and Discussion

### Systematic design to study Emajogi gene overexpression

Many of the genes encoded in phages are hypothetical and have no known homologs. We sought to understand the impact these phage genes have on *P. putida* growth using an overexpression screen. We separately cloned 47/50 genes from Emajogi into the pJLC140 plasmid backbone^13^ under an inducible arabinose promoter. We were unable to clone constructs for three genes (*gp5, gp14, gp33*). To assay cytotoxicity for each gene, we grew *P. putida* carrying each plasmid under four concentrations of arabinose (0%, 0.02%, 0.2%, and 2%) in two different media, LB and M9. A control plasmid expressing mCherry was used to normalize growth and expression. This plasmid does have low levels of leaky mCherry expression in the uninduced conditions (LB: 296 normalized relative fluorescent units (RFUs); M9: 61 normalized RFUs) and the highest level of mCherry expression was observed with 0.2% arabinose induction (**Supplemental Figure 1)**. The growth rate of each strain was measured in liquid culture using optical density and each gene was scored for cytotoxicity using a previously establish scoring system^4^. Genes with no impact on host growth when compared to the mCherry control at the same time point are scored as 0. Genes with a 1-3 log_2_ fold change compared to the mCherry control are score as 2, and genes with >3 log_2_ fold change in growth compared to mCherry are given a score of 3.

### Overexpression screen reveals eight genes that impact *P. putida* growth

In LB, only two genes were scored as cytotoxic. These were *gp18*, a predicted hypothetical gene, and *gp45*, a tail fiber gene (**Figure 1b**; **Supplemental Table 1**). We note that these genes are highly cytotoxic even without adding arabinose due to the low levels of leaky expression. We then turned to assessing cytotoxicity in M9 media, which can aid in a more controlled experiments due to the precisely defined media composition allowing for controlled cell growth and lower background for spectrophotometer assays. We found eight genes that negatively impacted *P. putida* cell growth in M9 (**Figure 1c**; **Supplemental Table 1**). These include the same two genes identified in LB with the same high level of cytotoxicity across all induction conditions. The remaining five genes were only cytotoxic at higher levels of induction. The RNA polymerase inhibitor gene (*gp17*) had 1.2-fold log_2_ reduction and 1.5-fold log_2_ reduction of growth with 0.2% and 2% arabinose induction, respectively. The remaining genes only showed growth reduction with 2% arabinose induction, these were *gp22*, a nucleotidyltransferase (4.2-fold); *gp23*, a DNA primase/helicase (1.2-fold); *gp30*, RNase H (1.2-fold); and *gp39*, a tail protein (3.8-fold).

Overall, gene cytotoxicity was only reported for 17% of Emajogi genes, which is lower than other phages that have been tested with similar assays (~25%)^4-6,14^. This lower rate could be due to the niche the phages were isolated from and how often they are naturally interacting with their host in these environment. Three genes were unable to be cloned, which may indicate they also have some cytotoxic effects.

In addition to providing an understanding of how these genes impact *P. putida* growth, this type of study can also provide insights on gene function in these phages. Six of the genes that were cytotoxic had annotated functions. Two of these were tail proteins, which likely have mechanisms to lyse the cell or arrest cell growth. The other two genes (*gp22* and *gp23*) are predicted to produce proteins that aid the phage in transcription and translation. Nucleotidyltransferase proteins have been shown to be components of toxin-antitoxin families and can alter tRNA molecules to arrest growth^15,16^. Similarly, overexpression of DNA primase/helicase, RNA polymerase inhibitor, and RNase H can disrupt normal DNA replication processes causing DNA damage responses to trigger^17^. The cytotoxicity is only observed at high levels of induction suggesting overexpression of these proteins disrupts normal transcription and translation processes.

The remaining cytotoxic gene, *gp18*, was predicted to encode a hypothetical protein. Additional computational analysis of gp18 was performed to identify any features associated with these proteins. No transmembrane domains were observed, no domains were found, and no significant homology across fifteen databases was found for this protein.

### Emajogi cytotoxic proteins are conserved across *Pseudomonas* phages

We sought to identify any homologs of the cytotoxic proteins in other known phages. The DNA primase/helicase (*gp23*), RNase H (*gp30*), and one of the tail proteins (*gp39*) had over 100 phage protein families that were near identical matches, suggesting these genes are conserved across bacteriophages. There were 61 homologs of the predicted nucleotidyltransferase (*gp22*) which included phages that infect *Dickeya, Xanthomonas*, and *Salmonella* (**Supplemental Table 1**). This protein is annotated in some of these phages as an Gabija anti-defense Gad2 family protein. This suggests that these proteins have conserved roles in transcription or translation.

The remaining four genes (*gp4, gp18, gp17*, and *gp45*) had lower numbers of homologs found in the databases. Overall, these four proteins had homologs in 60 other phages, with *gp4* identified in a mostly non-overlapping set of phages (**Supplemental Table 1**). While most of these were *Pseudomonas* phages, mostly infecting *Pseudomonas syringae*, we did identify homologs in phages that infect *Dickeya, Xanthomonas*, and *Salmonella*. The second tail fiber protein (*gp45*) which is cytotoxic in all conditions tested was found in 14 other phages, including *Dickeya* phage BIM BV-99 (**Supplemental Table 1**). This is not surprising since tail proteins have frequently been identified in other phage cytotoxicity screens and typically have some lysis capabilities to aid in cell entry^14^. The cytotoxic hypothetical proteins did have numerous homologs across multiple phages, which would suggest there is a conserved functional role for these proteins. We sought to further understand *gp18*, which was cytotoxic in all conditions tested.

### Transcriptional profiling of gp18 overexpression reveals a unique oxidative stress signature

To further explore the impacts of gp18, transcriptomic analysis was performed on an early exponential phase culture overexpressing gp18 with 0.2% arabinose. Differential expression profiling revealed significant shifts between the gp18 overexpression strain and either control (an empty vector and the inducible mCherry plasmid) (**Figure 2a**). More than 400 genes were significantly upregulated when gp18 was overexpressed (**Figure 2b; Supplemental Table 2)**. These genes include multiple transporter proteins, genes involved in cellular metabolism and oxidative stress, and prophage genes.

**Figure 2.**
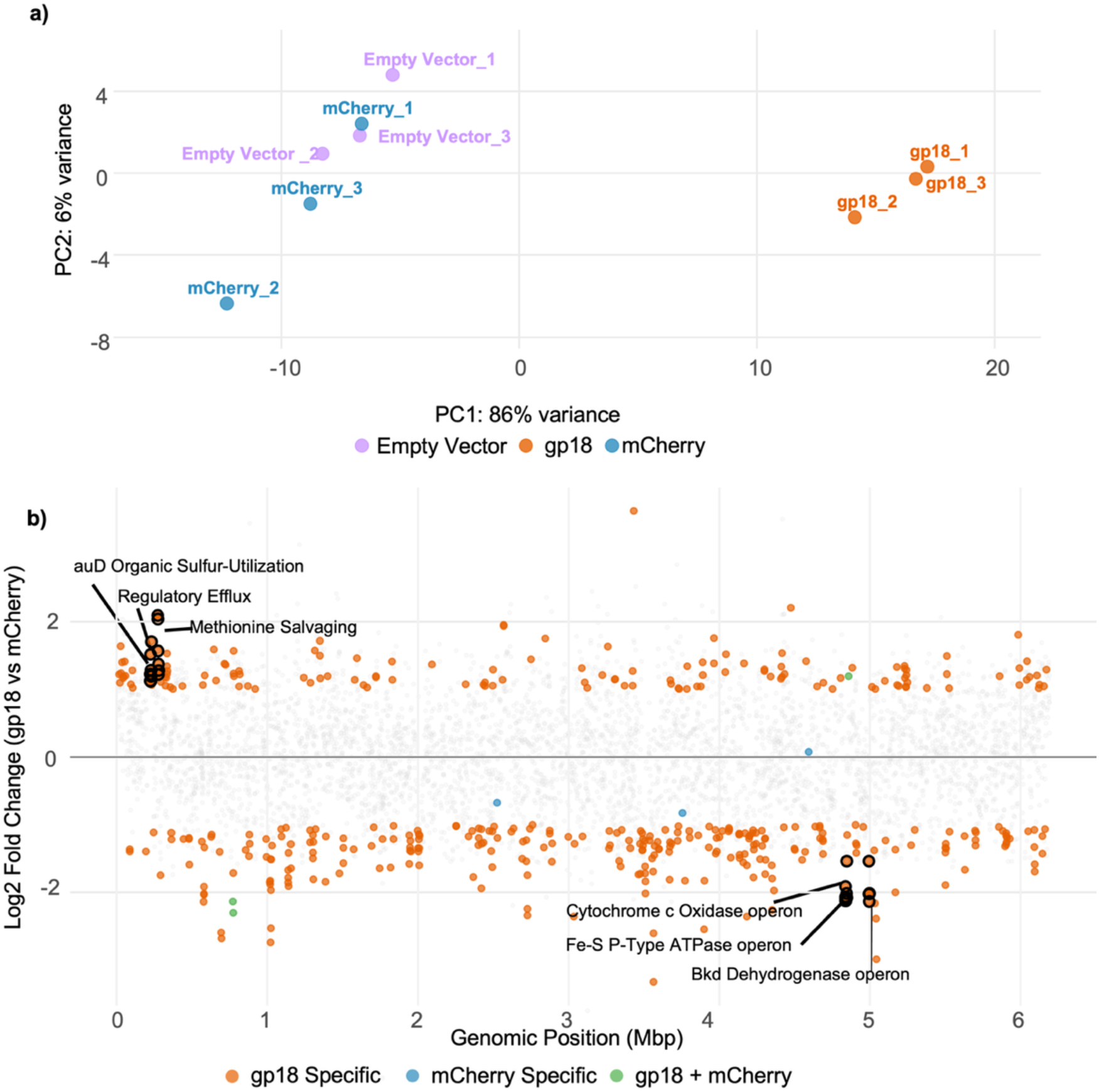
Transcriptional profile of gp18 overexpression in *P. putida* KT2440 reveals globally shifted transcriptome. **a)** Principal component analysis of the three transcriptomic samples. Each sample type is a triplicate of a culture overexpressing mCherry, gp18, or an empty vector. Gp18 has a distinct cluster of the three replicates and is distinct from the control samples. **b)** Log_2_ fold change values (gp18 compared to mCherry) for all genes are shown across the *P. putida* KT2440 genome (x-axis). Orange circles represent genes that are upregulated or down regulated in response to gp18 overexpression. Blue circles represent genes that were differentially expressed in response to mCherry overexpression. The single green circle represents the gene that was upregulated when either mCherry or gp18 were overexpressed but not the empty vector. The genes highlighted represent the most significantly upregulated and downregulated operons.

We chose to examine operons that were differentially expressed in the same direction (up or downregulated) in response to gp18 overexpression to reduce false positives and understand functional impacts of this protein. The three most upregulated operons based on average adjusted p-value were 1) a sulfur uptake and utilization locus (PP_0169–PP_0172), 2) a putative transcriptional regulator and efflux channel (PP_0175–PP_0177), and 3) a co-located methionine transport system and monooxygenase cluster associated with sulfur acquisition (PP_0219–PP_0221; PP_0223-PP_0224). These three operons would suggest there is a coordinated metabolic shift or overall cell-wide stress response. Conversely, expression of gp18 forced a coordinated downregulation of major stress response complexes, including 1) the 5-gene formate dehydrogenase-O (PP_0490-PP_0495), 2) the branched chain α-keto acid dehydrogenase complex (PP_4401-PP_4404), and 3) a cbb3-type 1 cytochrome c oxidase complex (PP_4250-PP_4253). From this data, we concluded that gp18 is involved in host take over and aids in shifting the cellular metabolism to viral production. This is because we see strong transcriptional changes in multiple genes and operons related to cellular metabolism and oxidative stress. It has been previously shown that changes in oxidative stress genes correlate with viral reprogramming to produce viral particles^18,19^. Further, many receptors and cell membrane proteins are differentially regulated suggesting a remodeling of the cell surface in response to overexpression of this protein, supporting a role in host cell take over.

## Conclusion

Here we describe the first overexpression screen of a *Pseudomonas* putida phage. Eight genes were cytotoxic in our screen, which is lower than average levels of cytotoxicity compared to other studies. Half of these genes were annotated to encode proteins that interact with DNA during replication and transcription. Two of these genes encoded tail proteins and the final two encoded hypothetical protein. Many homologs of the cytotoxic genes were identified across *Pseudomonas* phages, suggesting they may have generalize roles in phage production. Lastly, transcriptomic profiling of one of the hypothetical proteins, gp18, revealed a role for gp18 in cellular takeover of the host to convert the cell to viral production. Overall, overexpression screens are a useful tool to understand how phage genes impact their host and probe deeper into the viral “dark matter”.

## Materials and Methods

### Phage Gene Library Construction

Open reading frames (ORFs) encompassing the complete genomes of Emajogi (NCBI accession number PP4946445.1), were PCR amplified from 1 µl of filtered high titer Emajogi lysate using Invitrogen Platinum SuperFi II DNA Polymerase. Some genes could not be amplified via PCR and were ordered from Twist Biosciences. Each gene and the mCherry control was separately cloned into pJLC140^13^ using Gibson assembly and transformed into *E. coli* DH5alpha. Plasmids were verified by colony PCR using NEB Quikload Polymerase and the following primer pair that anneals to the pJLC140 plasmid surrounding the insert: 5’TGCTATGCCATAGCATTTTTATCC3’(AraBAD promoter) and 5’ACGCAGAAAGGCCCACC3’(BBa_B0014 double terminator). After sequence verification, plasmids were transformed into *P. putida* KT2440 using electroporation. Any plasmid that contained a gene with a positive score for cytotoxicity was confirmed via whole plasmid sequencing by Plasmidsaurs using Oxford Nanopore Technology with custom analysis and annotation.

### Kinetic Growth Assays and Induction Layout

*P. putida* KT2440_pE plasmids (**Supplemental Table 3**) were grown shaking overnight in LB at 30ºC, back diluted 1:100 into a 384-well plate and induced with one of the following L-arabinose concentrations 0%, 0.02%, 0.2%, or 2% in LB or M9 media. Optical density (OD_600_) and mCherry excitation/emissions (587nm/610nm) were measured on a Tecan Spark every ten minutes for 16-18 hours shaking at 30ºC. Each plate had triplicate samples for each media and induction concentration and an mCherry control.

### Calculation of RFUs for expression analysis

Normalized RFUs were calculated for the mCherry plasmid to determine the level of leaky expression. The triplicate RFUs for the mCherry plasmid were averaged for each time point with each inducer concentration. All wells from the same run that did not contain a mCherry plasmid were averaged to obtain a RFU for background.

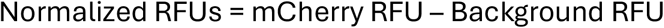

### Gene cytotoxicity calculation

The gene cytotoxicity score was calculated by averaging the OD_600_ of the three replicates of each gene at each concentration of arabinose in either LB or M9 at the final time point taken for each run (>16 hours). The OD_600_ was averaged for the final time point of mCherry expression in each run. The log_2_ fold change was then calculated as follows:

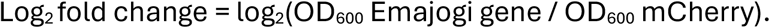

Cytotoxicity was scored as follows: 0: no impact on growth, 2: 1-3 log_2_ fold change, 3: >3 log_2_ fold change.

### Gp18 protein functional annotation

The protein sequence of gp18 (WYW02840.1) was used to determine a functional role for gp18. The protein sequence was input into a web-based InterPro^20^ search with standard parameters to identify any domains. The protein sequence was input into web-based deepTMHMM^21^ to identify any transmembrane domains. Lastly, the protein sequences was search against the standard databases included with DiMER^22^.

### Protein homolog search

The protein sequence of each cytotoxic gene was downloaded from NCBI and a web-based blastP^23^ search was used to determine any homologs in the clustered_nr database (accessed August 2026). Any protein that has >32.5% amino acid identity and e-values <10^-50^ were retained based on phamily criteria previously shown for mycobacteriophages^24^.

### RNA Isolation

*P. putida* KT2440 cultures harboring plasmids expressing either gp18, mCherry, or an empty vector backbone were induced with 0.2% arabinose and grown at 30ºC until they reached early log phase. Total RNA was extracted from these cultures using the Zymo QuickRNA Fungal/Bacterial kit. Sequencing libraries were prepared and sequenced using the Illumina NovaSeq platform at SeqCoast Genomics (Portsmouth, NH, USA).

### Differential Expression Analysis

Raw sequencing reads were aligned to the reference genome of *P. putida* KT2440 (NCBI accession: AE015451.2), and a genomic features count matrix was generated.

Downstream statistical analysis was performed in R (v4.4.0) using the DESeq2 package (v1.44.0)^25^. To control for the metabolic and translational burden associated with heterologous protein overexpression, the mCherry expression group was designated as the baseline reference level. Contrasts were established to isolate gp18-specific transcriptional shifts relative to the mCherry control, filtering out general vector and transcription-related stress. A secondary contrast between the mCherry control and the empty plasmid group was utilized as a control filter to ensure the neutrality of the expression baseline. Replicate quality and distribution were assessed via Principal Component Analysis (PCA) and dispersion trend modeling.

The following criteria were used to identify operons that were significantly changed. A genomic window was classified as an actively co-regulated operon if it fulfilled two stringent criteria: 1) A minimum of three genes on the same strand transcriptionally shifting in the same direction (upregulated or downregulated). One gene not in this uniform direction was permitted per operon. 2) The transcriptional change of each gene within the operon had an absolute fold change of more than 2 and an adjusted *p*-value of <0.01.

## Acknowledgments

We thank Jesse Cahill for sharing the pJLC140 expression plasmid backbone used in this study. We thank Hedvig Tamman for the Emajogi phage. We thank Lillian Lowrey for internal review of this manuscript.

Research funding provided by Laboratory Directed Research and Development program at Sandia National Laboratories (233066 to CMM).

Sandia National Laboratories is a multimission laboratory managed and operated by National Technology & Engineering Solutions of Sandia, LLC, a wholly owned subsidiary of Honeywell International Inc., for the U.S. Department of Energy’s National Nuclear Security Administration under contract DE-NA0003525. This paper describes objective technical results and analysis.

Any subjective views or opinions that might be expressed in the paper do not necessarily represent the views of the U.S. Department of Energy or the United States Government. This article has been authored by an employee of National Technology & Engineering Solutions of Sandia, LLC under Contract No. DE-NA0003525 with the U.S. Department of Energy (DOE). The employee owns all right, title and interest in and to the article and is solely responsible for its contents. The United States Government retains and the publisher, by accepting the article for publication, acknowledges that the United States Government retains a non-exclusive, paid-up, irrevocable, world-wide license to publish or reproduce the published form of this article or allow others to do so, for United States Government purposes. The DOE will provide public access to these results of federally sponsored research in accordance with the DOE Public Access Plan https://www.energy.gov/downloads/doe-public-access-plan.

## Data Availability

We used the following phage genome accession number for Emajogi PP496445. All plasmids are listed in **Supplementary Table 3**. The raw RNA sequencing reads are at NCBI under BioProject PRJNA1529819.

