## Supplemental Figure for "Genome-wide overexpression screen of *Pseudomonas putida* phage Emajogi"

**Supplemental Material**

### Supplemental Tables

**Supplemental Table 1. Emajogi gene list with coordinates, function, cytotoxicity score across media types, and identified homologs.**

**Supplemental Table 2. Significantly differentially expressed genes in *P. putida* KT2440 in response to overexpression of gp18.** All genes that were up- or down-regulated in response to Emajogi gp18 overexpression are shown. The $log_2$ fold change (fc) was calculated for Emajogi specific effect and plasmid effect. There is a category in the final column that identifies which dataset it is considered part of.

**Supplemental Table 3. Plasmids generated in this study.**

### Supplemental Figures

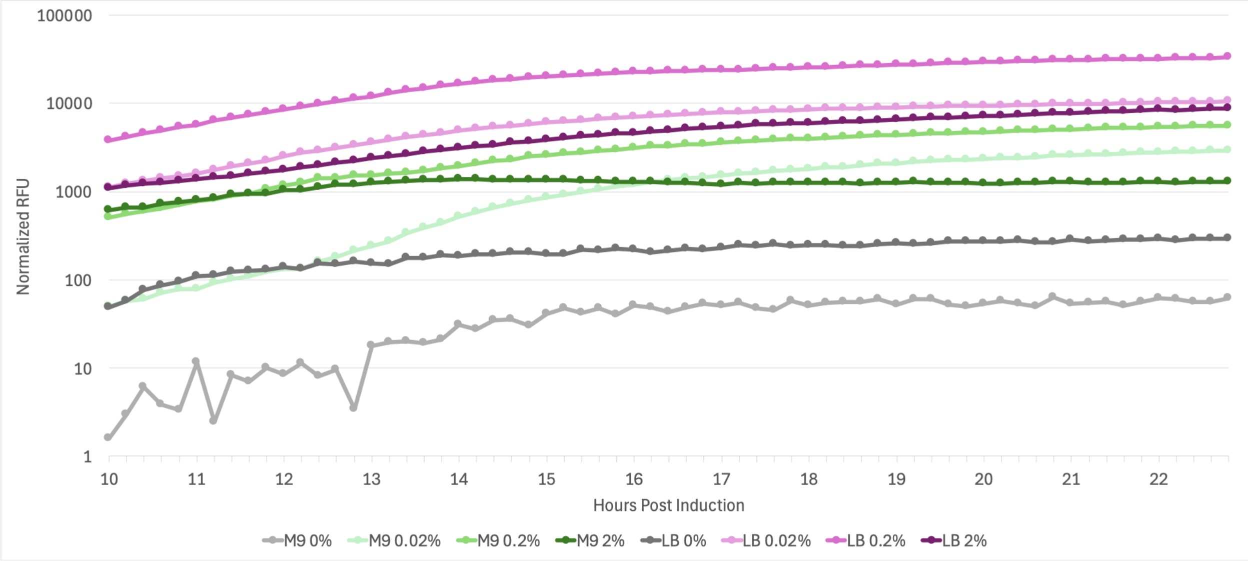


**Supplementary Figure 1. RFUs of mCherry control plasmid induced with arabinose in LB and M9.** Normalized RFU values are shown on a logarithmic y-axis. The hours post arabinose induction are shown on the x-axis. The normalized RFU values are averages of three triplicates with background RFUs removed from the same experimental plate. We chose to show time points after 10-hours of growth to aid in visualization. Induction in M9 media is shown in green, induction in LB media is shown in pink, and uninduced are shown in grey.
